# CityNet - Deep Learning Tools for Urban Ecoacoustic Assessment

**DOI:** 10.1101/248708

**Authors:** A. J. Fairbrass, M. Firman, C. Williams, G. J. Brostow, H. Titheridge, K. E. Jones

## Abstract

1. Cities support unique and valuable ecological communities, but understanding urban wildlife is limited due to the difficulties of assessing biodiversity. Ecoacoustic surveying is a useful way of assessing habitats, where biotic sound measured from audio recordings is used as a proxy for biodiversity. However, existing algorithms for measuring biotic sound have been shown to be biased by non-biotic sounds in recordings, typical of urban environments.
2. We develop CityNet, a deep learning system using convolutional neural networks (CNNs), to measure audible biotic (CityBioNet) and anthropogenic (CityAnthroNet) acoustic activity in cities. The CNNs were trained on a large dataset of annotated audio recordings collected across Greater London, UK. Using a held-out test dataset, we compare the precision and recall of CityBioNet and CityAnthroNet separately to the best available alternative algorithms: four acoustic indices (AIs): Acoustic Complexity Index, Acoustic Diversity Index, Bioacoustic Index, and Normalised Difference Soundscape Index, and a state-of-the-art bird call detection CNN (bulbul). We also compare the effect of non-biotic sounds on the predictions of CityBioNet and bulbul. Finally we apply CityNet to describe acoustic patterns of the urban soundscape in two sites along an urbanisation gradient.
3. CityBioNet was the best performing algorithm for measuring biotic activity in terms of precision and recall, followed by bulbul, while the AIs performed worst. CityAnthroNet outperformed the Normalised Difference Soundscape Index, but by a smaller margin than CityBioNet achieved against the competing algorithms. The CityBioNet predictions were impacted by mechanical sounds, whereas air traffic and wind sounds influenced the bulbul predictions. Across an urbanisation gradient, we show that CityNet produced realistic daily patterns of biotic and anthropogenic acoustic activity from real-world urban audio data.
4. Using CityNet, it is possible to automatically measure biotic and anthropogenic acoustic activity in cities from audio recordings. If embedded within an autonomous sensing system, CityNet could produce environmental data for cites at large-scales and facilitate investigation of the impacts of anthropogenic activities on wildlife. The algorithms, code and pre-trained models are made freely available in combination with two expert-annotated urban audio datasets to facilitate automated environmental surveillance in cities.

## INTRODUCTION

Over half of the world’s human population now live in cities (UN-DESA 2016) and urban biodiversity can provide people with a multitude of health and well-being benefits including improved physical and psychological health (Natural England 2016; Crouse *et al*. 2017). Cities can support high biodiversity including native endemic species (Aronson *et al*. 2014), and act as refuges for biodiversity that can no longer persist in intensely managed agricultural landscapes surrounding cities (Hall *et al*. 2016). However, our understanding of urban biodiversity remains limited (Faeth, Bang & Saari 2011; Beninde, Veith & Hochkirch 2015). One reason for this is the difficulties associated with biodiversity assessment, such as gaining repeated access to survey sites and the resource intensity of traditional methods (Farinha-Marques *et al*. 2011). This inhibits our ability to conduct the large-scale assessment that is necessary for understanding urban ecosystems.

Ecoacoustic surveying has emerged as a useful method of large-scale quantification of ecological communities and their habitats (Sueur & Farina 2015). Passive acoustic recording equipment facilitates the collection of audio data over long time periods and large spatial scales with fewer resources than traditional survey methods (Digby *et al.* 2013). A number of automated methods have been developed to measure biotic sound in the large volumes of acoustic data that are typically produced by ecoacoustic surveying (Sueur & Farina 2015). For example, Acoustic Indices (AIs) use the spectral and temporal characteristics of acoustic energy in sound recordings to produce whole community measures of biotic and anthropogenic sound (Sueur *et al.* 2014). However, several commonly used AIs have been shown to be biased by non-biotic sounds (Towsey *et al.* 2014; Fuller *et al.* 2015; Gasc *et al.* 2015a), and are not suitable for use in the urban environment without the prior removal of certain non-biotic sounds from recordings (Fairbrass *et al.* 2017).

Machine learning (ML) is being increasingly applied to biodiversity assessment and monitoring because it facilitates the detection and classification of ecoacoustic signals in audio data (Acevedo *et al.* 2009; Walters *et al.* 2012; Stowell & Plumbley 2014). Using annotated audio datasets of soniferous species, a ML model can be trained to recognise biotic sounds based on multiple acoustic characteristics, or features, and to associate these features with taxonomic classifications, and can then assign a probabilistic classification to sounds within recordings. AIs only use a limited number of acoustic features in their calculations, such as spectral entropy within defined frequency bands (Boelman *et al.* 2007; Villanueva-Rivera *et al.* 2011; Kasten *et al.* 2012) or entropy changes over time (Pieretti, Farina & Morri 2011). Additionally, the relationship between the features and the algorithm outputs are chosen by a human, rather than learned automatically from an annotated dataset. In contrast, ML algorithms can utilise many more features in their calculations, and the relationship between inputs and outputs is determined automatically based on the annotated training data provided. Convolutional Neural Networks, CNNs (or Deep learning) (LeCun, Bengio & Hinton 2015) can even choose, based on the annotations in the training dataset, the features that best discriminate different classes in datasets without being specified a priori, and can take advantage of large quantities of training data where their ability to outperform human defined algorithms increases as more labelled data become available.

Species-specific ML algorithms have been developed to automatically identify the sounds emitted by a range of soniferous organisms including birds (Stowell & Plumbley 2014), bats (Walters *et al.* 2012; Zamora-Gutierrez *et al*. 2016), amphibians (Acevedo *et al.* 2009) and invertebrates (Chesmore & Ohya 2004). However, these algorithms are focussed on a small number of species limiting their usefulness for broad classification tasks across communities. More recently, algorithms that detect whole taxonomic groups are being developed, for example bird sounds in audio recordings from the UK and the Chernobyl Exclusion Zone (Grill & Schlüter 2017), but these algorithms remain untested on noisy audio data from urban environments. There are currently no algorithms that produce whole community measures of biotic sound that are known to be suitable for use in acoustically complex urban environments.

Here, we develop the CityNet acoustic analysis system, which uses two CNNs for measuring audible (0-12 kHz) biotic (CityBioNet) and anthropogenic (CityAnthroNet) acoustic activity in audio recordings from urban environments. We use this frequency range as it contains the majority of sounds emitted by audible soniferous species in the urban environment (Fairbrass *et al.* 2017). The CNNs were trained using CitySounds2017, an expert-annotated dataset of urban sounds collected across Greater London, UK that we develop here. We compared the performance of CityNet using a held-out dataset by comparing the algorithms’ precision and recall to four commonly used AIs: Acoustic Complexity Index (ACI) (Pieretti, Farina & Morri 2011), Acoustic Diversity Index (ADI) (Villanueva-Rivera *et al.* 2011), Bioacoustic Index (BI) (Boelman *et al.* 2007), Normalised Difference Soundscape Index (NDSI) (Kasten *et al.* 2012), and to bulbul, a state-of-the-art algorithm for detecting bird sounds in order to summarise avian acoustic activity (Grill & Schlüter 2017). As the main focus of the study was the development of algorithms for ecoacoustic assessment of biodiversity in cities, we conducted further analysis on the two best performing algorithms for measuring biotic sound, CityBioNet and bulbul, by investigating the effect of non-biotic sounds on the accuracy of the algorithms. Finally, we applied CityNet to investigate daily patterns of biotic and anthropogenic sound in the urban soundscape.

## MATERIALS AND METHODS

We developed two CNN models, CityBioNet and CityAnthroNet within the CityNet system to generate measures of biotic and anthropogenic sound, respectively. The CityNet pipeline (Figure 1) consisted of 7 main steps as follows:

1. *Record audio:* Audible frequency (0-12 kHz) *.wav* audio recordings were made using a passive acoustic recorder.
2. *Audio conversion to Mel spectrogram:* Each audio file was automatically converted to a Mel spectrogram representation with 32 frequency bins, represented as rows in the spectrogram, using a temporal resolution of 21 columns per second of raw audio. Before use in the classifier, each spectrogram *S* was converted to a log-scale representation, using the formula log(A + B * S). For biotic sound detection the parameters A = 0.001 and B = 10.0 were used, while for anthropogenic sound detection the parameters A = 0.025 and B = 2.0 were used.
3. *Extract window from spectrogram:* A single input to the CNN comprised a short spectrogram chunk *Ws*, 21 columns in width, representing 1 second of audio.
4. *Apply different normalisation strategies:* There are many different methods for preprocessing spectrograms before they are used in ML; for example whitening (Lee *et al.* 2009) and subtraction of mean values along each frequency bin (Aide *et al.* 2013). CNNs are able to accept inputs with multiple channels of data, for example the red, green and blue channels of a colour image. We exploited the multiple input channel capability of our CNN by providing as input four spectrograms each pre-processed using a different normalisation strategy (see Supplementary Methods), which gave considerable improvements to network accuracy above any single normalisation scheme in isolation. After applying different normalisation strategies, the input to the network consisted of a 32 x 21 x 4 tensor.
5. *Apply CNN classifier:* As described above, classification was performed with a CNN, whose parameters were learnt from training data. The CNN comprised a series of layers, each of which modified its input data with parameterised mathematical operations which were optimised to improve classification performance during training (see Supplementary Methods for details). The final layer produced the prediction of presence or absence of biotic or anthropogenic sound.
6. *Make prediction for each moment in time:* At test time, steps (3)-(5) were repeated every 1 second throughout the audio file, to give a measure of biotic or anthropogenic activity throughout time. Predictions for each chunk of audio were made independently.
7. *Summarise:* Where appropriate, the chunk-level predictions were summarised to gain insights into trends over time and space. For example, predicted activity levels for each half-hour window could be averaged to inspect the level of biotic and anthropogenic activity at different times of day.

**Figure 1.**
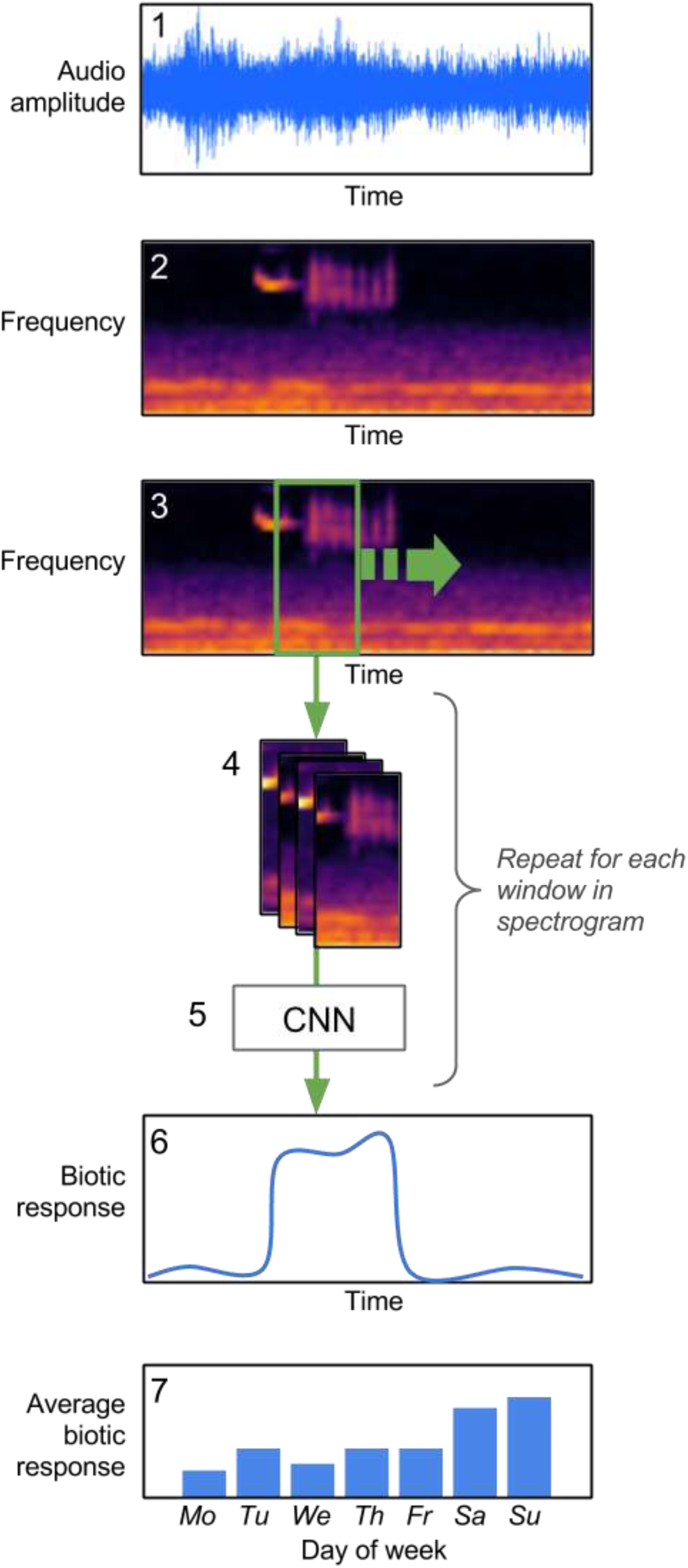
The CityNet analysis pipeline for measuring biotic and anthropogenic acoustic activity. Raw audio (1), recorded in the field, is converted to a spectrogram representation (2). A sliding window is run across the time dimension, and a window of the spectrogram extracted at each step (3). This spectrogram window is pre-processed with four different normalisation strategies, and the results concatenated. This stack of spectrograms is passed through a CNN (5), which was trained on CitySounds2017_*train*_. The CNN gives, at each 1-second time step, a prediction of the presence/absence of biotic or anthropogenic acoustic activity (6). Finally, these per-time-step measures can be aggregated to give summaries over time or space (7).

The ML pipeline was written in Python v.2.7.12 (Python Software Foundation 2016) using Theano v.0.9.0 (The Theano Development Team *et al*. 2016) and Lasagne v.0.2 (Dieleman *et al.* 2015) for ML and librosa v.0.4.2 (McFee *et al.* 2015) for audio processing.

### Acoustic Dataset

We selected 63 green infrastructure (GI) sites in and around Greater London, UK to collect audio data to train and test the CityNet algorithms. These sites represent a range of GI in and around Greater London in terms of GI type, size and urban intensity. Each site was sampled for 7 consecutive days systematically across the months of May to October between 2013 and 2015 (Figure 2, Table S1). At each location, a Song Meter SM2+ digital audio field sensor (Wildlife Acoustics, Inc., Concord, Massachusetts, USA) was deployed, recording sound between 0 and 12 kHz at a 24 kHz sample rate. The sensor was equipped with a single omnidirectional microphone (frequency response: ‐35±4 dB) oriented horizontally at a height of 1m. Files were saved in *.wav* format onto a SD card. Audio was recorded in computationally manageable chunks of 29 minutes of every 30 mins (23.2 hours of recording per day), which were divided into 1-minute audio files using Slice Audio File Splitter (NCH Software Inc. 2014), leading to a total of 613,872 discrete minutes of audio recording (9,744 minutes for each of the 63 sites). This constituted the CitySounds2017 dataset.

**Figure 2.**
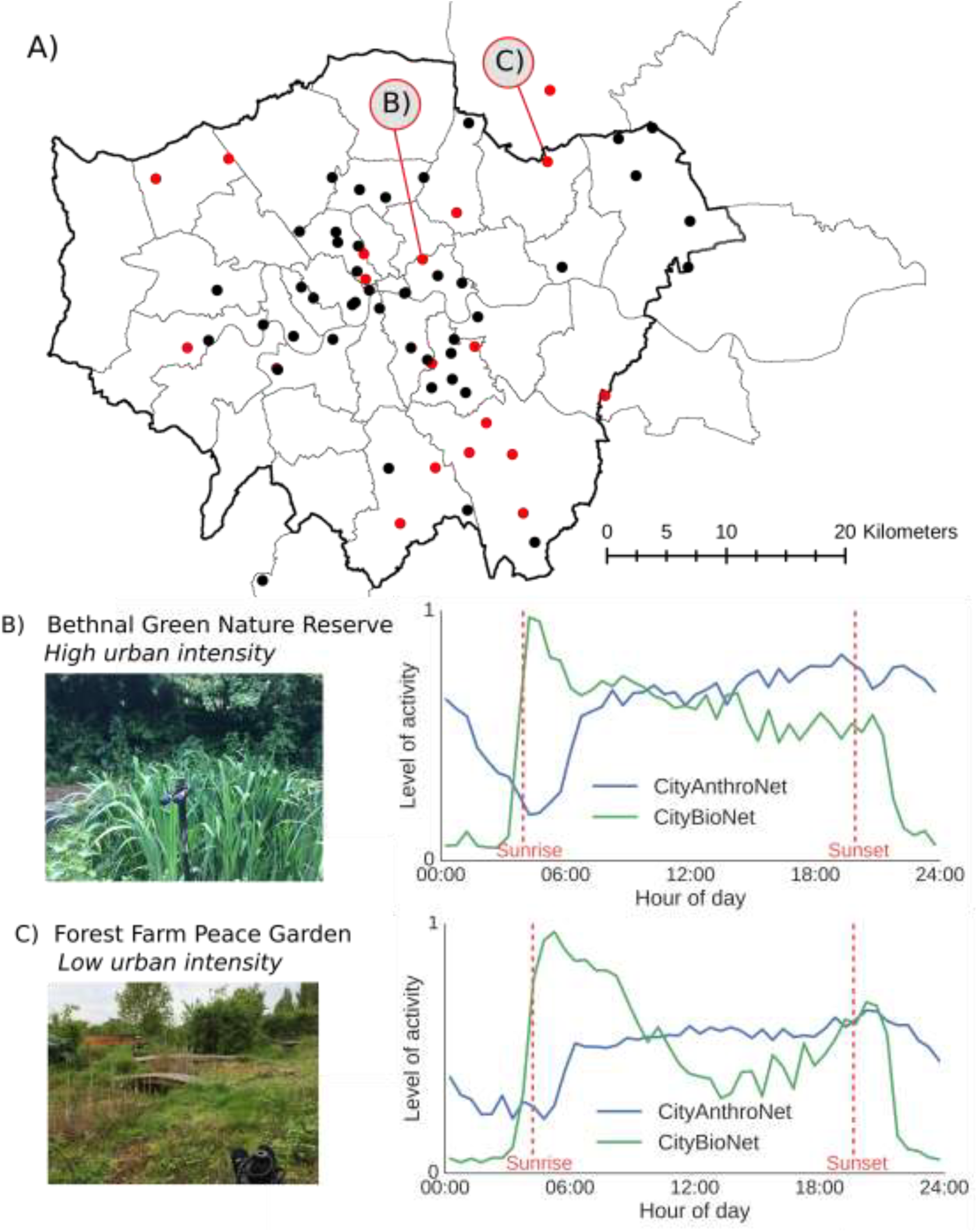
Location of study sites and average daily acoustic patterns at two sites along an urbanisation gradient. Points in (A) represent locations used for the training dataset, CitySounds2017_*train*_ (black) and testing dataset, CitySounds2017_*test*_ (red). Here CityNet was run across the entire 7 days of recording at two sites of high (B) and low (C) urban intensity to predict the presence/absence of biotic and anthropogenic sound at each second of the week using a 0.5 probability threshold. The predicted number of seconds containing biotic and anthropogenic sound for each half-hour period was averaged over the week to produce average daily patterns of acoustic activity. Greater London boundary indicated with bold line. Boundary data from the UK Census (http://www.ons.gov.uk/, accessed 04/11/2014).

### Acoustic Training Dataset

To create our training dataset (CitySounds2017_*train*_) we randomly selected twenty five 1-minute recordings from 70% of the study sites (44 sites, 1100 recordings). A.F. manually annotated the spectrograms of each recording, computed as the log magnitude of a discrete Fourier transform (non-overlapping Hamming window size=720 samples=10 ms), using AudioTagger (available at https://github.com/groakat/AudioTagger). Spectrograms were annotated by localising the time and frequency bands of discrete sounds by drawing bounding boxes as tightly as visually possible within spectrograms displayed on a Dell UltraSharp 61cm LED monitor. Types of sound, such as “invertebrate”, “rain”, and “road traffic”, were identified by looking for typical patterns in spectrograms (Figure S1), and by listening to the audio samples represented in the annotated parts of the spectrogram. Categories of sounds were then grouped into biotic, anthropogenic and geophonic classes following Pijanowski *et al*. (2011), where we define biotic as sounds generated by non-human biotic organisms, anthropogenic as sounds associated with human activities, and geophonic as non-biological ambient sounds e.g. wind and rain.

### Acoustic Testing Dataset and Evaluation

To evaluate the performance of the CityNet algorithms, we created a testing dataset (CitySounds2017_*teit*_) by strategically selecting 40 recordings from CitySounds2017 from the remaining 30% of sites (19 sites) that contained a range of both biotic and anthropogenic acoustic activity. CitySounds2017_*test*_ was sampled from different recording sites to CitySounds2017_*train*_ to demonstrate that the CityNet algorithms generalise to sounds recorded at new site locations (Figure 2, Table S1). To optimise the quality of the annotations in CitySounds2017_*test*_, we selected five human labellers to separately annotate the sounds within the audio recordings (using the same methods as above) to create a single annotated test dataset. Conflicts were resolved using a majority rule, and in cases where there was no majority, we used our own judgement on the most suitable classification. Our CitySounds2017 annotated training and testing datasets are available at https://figshare.com/s/adab62c0591afaeafedd.

Using the CitySounds2017_*test*_ dataset, we separately assessed the performance of the two CityNet algorithms, CityBioNet and CityAnthroNet, using two measures: precision and recall. The CityBioNet and CityAnthroNet algorithms give a probabilistic estimate of the level of biotic or anthropogenic acoustic activity for each 1-second audio chunk as a number between 0 and 1. Different thresholds could be used to convert these probabilities into sound category assignments (e.g. ‘sound present’ or ‘sound absent’). At each threshold, a value of precision and recall was computed, where precision was the fraction of 1-second chunks correctly identified as containing the sound according to the annotations in CitySounds2017_*test*_, and recall was the fraction of 1-second chunks labelled as containing the sound which was retrieved by the algorithm under that threshold. As the threshold was swept between 0 and 1, the resulting values of precision and recall were plotted as a precision-recall curve. Summary statistics were computed for the average precision under all the threshold values and the recall when the threshold chosen gave a precision of 0.95. Using a threshold of 0.5 on the predictions, confusion matrices were calculated showing how each moment of time was classified relative to the annotations. These analyses were conducted in Python v.2.7.12 (Python Software Foundation 2016) using Scikit-learn v.0.18.1 (Pedregosa *et al.* 2011) and Matplotlib v.1.5.1 (Hunter 2007).

### Competing Algorithms

We also compared the precision and recall of the CityNet algorithms to acoustic measures produced by four AIs: Acoustic Complexity Index (ACI) (Pieretti, Farina & Morri 2011), Acoustic Diversity Index (ADI) (Villanueva-Rivera *et al.* 2011), Bioacoustic Index (BI) (Boelman *et al.* 2007), and Normalised Difference Soundscape Index (NDSI) (Kasten *et al.* 2012). The NDSI generates a measure of anthropogenic disturbance according to the formula

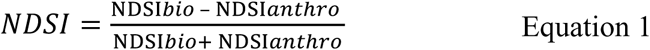

where NDSI_*bio*_ and NDSI_*anthro*_ are the total biotic and anthropogenic acoustic activity in each recording, respectively. Rather than compare CityNet to the NDSI, we compared the biotic (NDSI_*bio*_) and anthropogenic (NDSI_*anthro*_) elements of the NDSI to the measures produced by CityBioNet and CityAnthroNet, respectively, as these were more comparable. As the AIs are all designed to give a summary of acoustic activity for an entire file, they were analysed on the CitySounds2017_*test*_ dataset by treating each 1-second chunk of audio as a separate sound file to enable direct comparisons to CityNet. The AI measures do not have a natural threshold for classification into biotic/non-biotic sound, meaning we could not calculate confusion matrices. However, a threshold between their lowest value and their highest value was used in combination with the range of precision and recall values to form precision-recall curves. All AIs were calculated in R v.3.4.1 (R Core Team 2017) using the ‘seewave’ v.1.7.6 (Sueur, Aubin & Simonis 2008) and ‘soundecology’ v.1.2 (Villanueva-Rivera & Pijanowski 2014) packages.

The precision and recall of CityBioNet was also compared to bulbul (Grill & Schlüter 2017), an algorithm for detecting bird sounds in entire audio recordings in order to summarise avian acoustic activity which was the winning entry in the 2016-7 Bird Audio Detection challenge (Stowell *et al.* 2016). Like CityNet, bulbul is a CNN-based classifier which uses spectrograms as input. However, it does not use the same normalisation strategies as CityNet, and it was not trained on data from noisy, urban environments. Bulbul was applied to each second of audio data in CitySounds2017_*test*_, using the pre-trained model provided by the authors together with their code.

### Impact of Non-Biotic Sounds

We conducted additional analysis on the non-biotic sounds that affect the predictions of CityBioNet and bulbul, as these were found to be the best performing algorithms for measuring biotic sound. To do this, we created subsets of the CitySounds2017_*test*_ dataset comprising all the seconds that contained a range of non-biotic sounds, e.g. a road traffic data subset containing all of the seconds in CitySounds2017_*test*_ where the sound of road traffic was present. We then used a Chi-squared test to identify significant differences in the proportion of seconds in which the presence/absence of biotic sound at threshold 0.5 was correctly predicted in the full and subset datasets by each algorithm, and the Cramer’s V statistic was used to assess the effect size of differences (Cohen 1992). These analyses were conducted in R v.3.4.1 (R Core Team 2017).

### Ecological Application

We used CityNet to generate daily average patterns of biotic and anthropogenic acoustic activity for two study sites across an urbanisation gradient (sites E29RR and IG62XL with high and low urbanisation respectively, Table S1). To control for the date of recording; both sites were surveyed between May and June 2015. CityNet was run over the entire 7 days of recordings from each site to predict the presence/absence of biotic and anthropogenic sound for every 1-second audio chunk using a 0.5 probability threshold. Measures of biotic and anthropogenic activity were created for each half hour window between midnight and midnight by averaging the predicted number of seconds containing biotic or anthropogenic sound within that window over the entire week.

## RESULTS

### Acoustic Performance

CityBioNet had an average precision of 0.934 and recall of 0.710 at 0.95 precision, while CityAnthroNet had an average precision of 0.977 and recall of 0.858 at 0.95 precision (Table 1, Figure 3). In comparison the ACI, ADI, BI and NDSI_*bio*_ had a lower average precision (0.663, 0.439, 0.516, and 0.503, respectively) and lower recall at 0.95 (all less than 0.01). CityBioNet also outperformed bulbul which had an average precision of 0.872 and recall at 0.95 of 0.398 (Table 1). In comparison to CityAnthroNet, the NDSI_*anthro*_ had a lower average precision (0.975) and lower recall at 0.95 precision (0.815). When biotic sound was present in recordings, CityBioNet correctly predicted the presence of biotic sound (True Positives) in a greater proportion of audio data than bulbul (33.2% in comparison with 18.5%, for CityBioNet and bulbul respectively) (Figure 4). However, CityBioNet failed to correctly predict the presence of biotic sound (False Negatives) in 1.7% of recordings in comparison with 1.0% incorrect predictions by bulbul. When biotic sound was absent from recordings, CityBioNet correctly predicted the absence of biotic sound (True Negatives) in 51.6% of the audio data in comparison with 52.6% for bulbul, and CityBioNet failed to correctly predict the absence of biotic sound (False Positives) in 13.5% of audio data in comparison with 20.0% incorrect predictions by bulbul (Figure 4).

**Figure 3.**
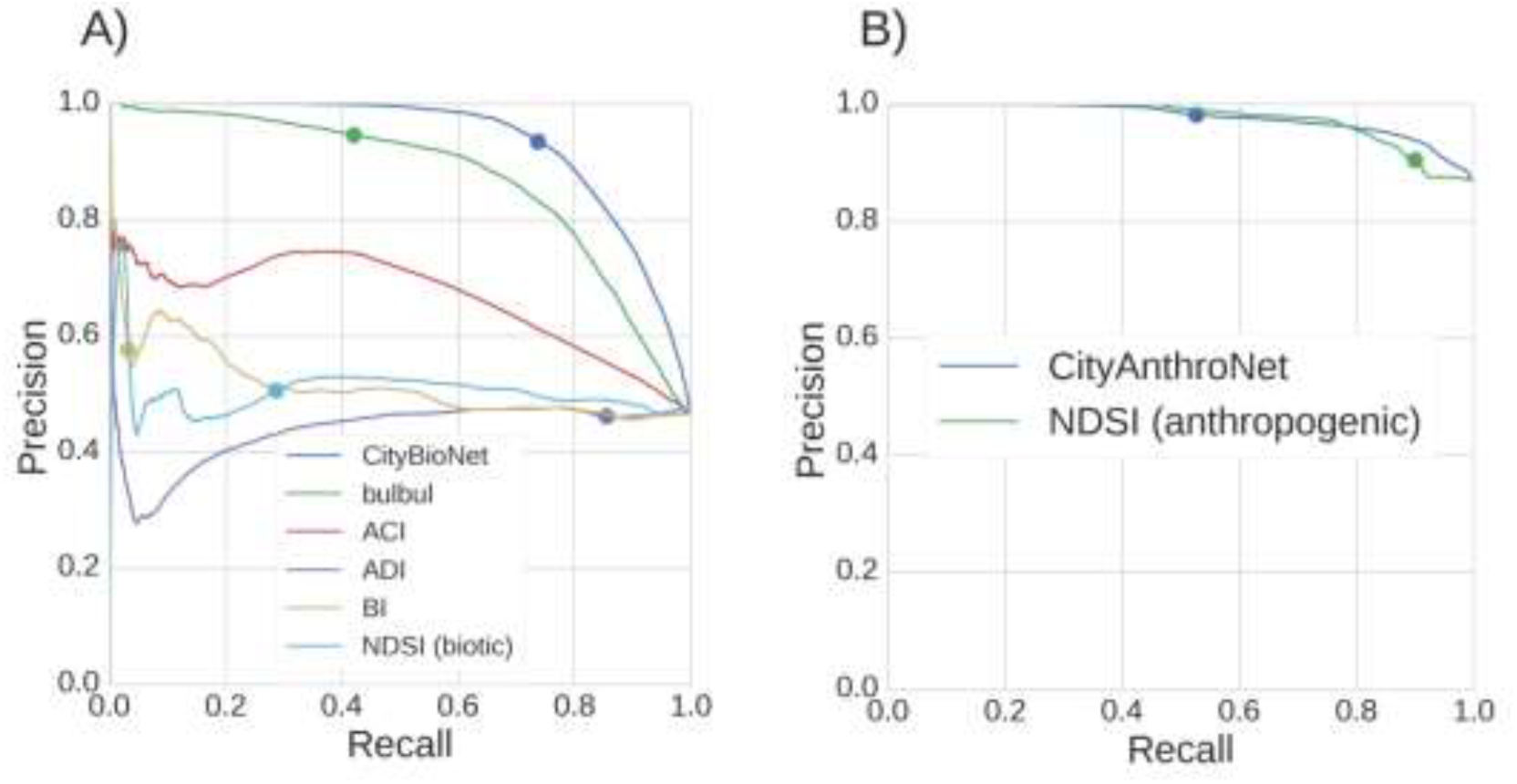
Precision-recall curves for CityNet and competing algorithms predicting A) biotic and B) anthropogenic acoustic activity for each 1-second audio chunk in the CitySounds2017_*test*_ dataset. Dots indicate the precision and recall values at a threshold value of 0.5. ACI represents Acoustic Complexity Index, ADI Acoustic Diversity Index, BI Bioacoustic Index, and NDSI_*bio*_ and NDSI_*anthro*_ biotic and anthropogenic Normalised Difference Soundscape Index, respectively.

**Table 1.** Average precision and recall results for CityNet and competing algorithms for each 1-second audio chunk in the CitySounds2017_*test*_ dataset. Recall results are presented at 0.95 precision. Higher values are better for both metrics. The highest values in each section are shown in bold. ACI represents Acoustic Complexity Index, ADI Acoustic Diversity Index, BI Bioacoustic Index, and NDSI_*bio*_ and NDSI_*anthro*_ biotic and anthropogenic Normalised Difference Soundscape Index, respectively.

| Acoustic Measures | Recall at 0.95 precision | Average precision |
| --- | --- | --- |
| ***Biotic*** |  |  |
| CityBioNet | **0.710** | **0.934** |
| Bulbul | 0.398 | 0.872 |
| ACI | 0.000 | 0.663 |
| ADI | 0.001 | 0.439 |
| BI | 0.002 | 0.516 |
| NDSI<sub>biotic</sub> | 0.000 | 0.503 |
| ***Anthropogenic*** |  |  |
| CityAnthroNet | **0.858** | **0.977** |
| NDSI<sub>anthro</sub> | 0.815 | 0.975 |

**Figure 4.**
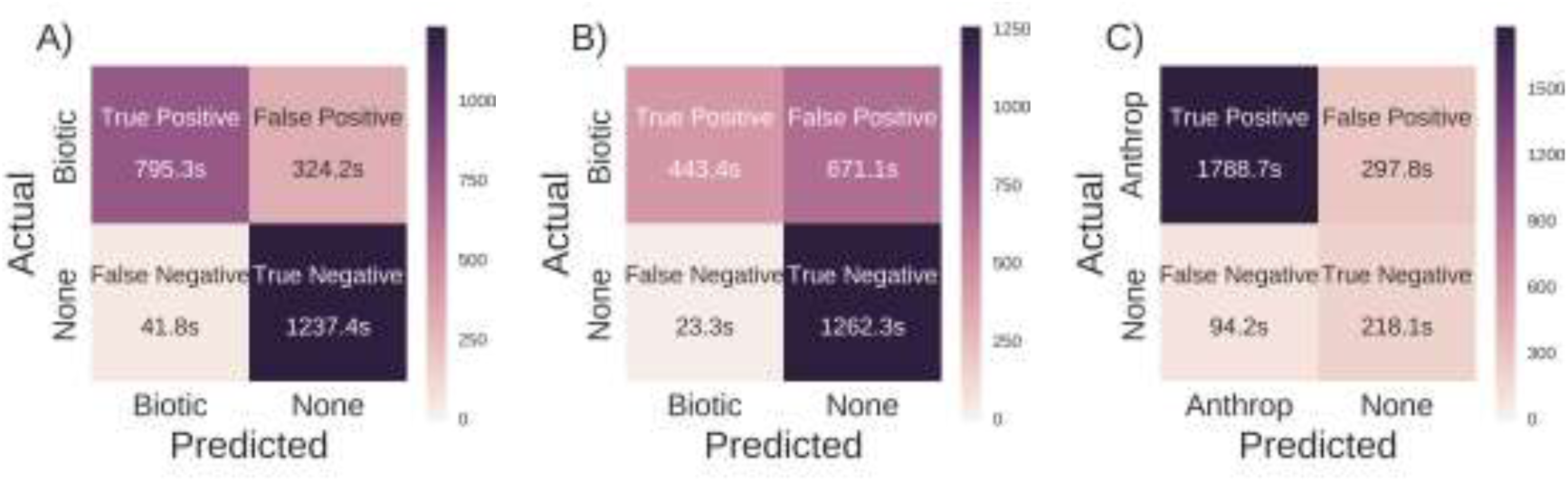
Confusion matrices comparing the predicted acoustic activity of A) CityBioNet, B), bulbul, and C) CityAnthroNet for each 1-second audio chunk in the CitySounds2017_*test*_ dataset. Numbers in each cell report the number of 1-second audio clips in the CitySounds2017_*test*_ dataset predicted either correctly (True Positives and True Negatives) or incorrectly (False Positives and False Negatives) as containing biotic (A and B) or anthropogenic (C) sound. To create the confusion matrices, the probabilistic predictions from the classifiers are converted to binary classifications using a threshold that gives a precision of 0.95.

### Impacts of Non-Biotic Sounds

CityBioNet was strongly (Cramer’s V effect size >0.5) negatively affected by mechanical sound (the presence/absence of biotic sound was correctly predicted in 28.60% less of the data when mechanical sounds were also present) (Table 2). Bulbul was moderately (Cramer’s V effect size 0.1-0.5) negatively affected by the sound of air traffic and wind (the presence/absence of biotic sound was correctly predicted in 5.34% and 6.93% less of the data when air traffic and wind sounds were also present in recordings, respectively).

**Table 2.** Impact of non-biotic sounds on the CityBioNet and bulbul predictions. Values represent differences in the proportion of 1-second audio chunks in the full CitySound2017_*test*_ dataset (40 minutes) and the subset datasets (size in time indicated in left-hand column) in which the presence/absence of biotic sound was correctly predicted by both algorithms, (chisquared test statistic for difference in proportions of successes in each dataset, and Cramer’s V effect size measure). Effect sizes indicated as <0.1 (*), 0.1-0.3 (**) and >0.5 (***).

| Sound Type | CityBioNet | Bulbul |
| --- | --- | --- |
| *Anthropogenic* |  |  |
| Air traffic (9m 4s) | -2.11 (30.35, 0.05)* | -5.34 (162.73, 0.12)** |
| Mechanical (11s) | -28.60 (134.38, 0.77)*** | 0.02 (0.01, 0.01)* |
| Road traffic (29m 15s) | 0.79 (10.15, 0.02)* | 1.41 (27.67, 0.03)* |
| Siren (1m 21s) | 2.28 (5.73, 0.06)* | 3.70 (12.95, 0.09)* |
| *Geophonic* |  |  |
| Rain (2m 44s) | -0.77 (1.29, 0.02)* | -1.51 (4.17, 0.04)* |
| Wind (53s) | 0.76 (0.47, 0.02)* | -6.93 (33.11, 0.17)** |

### Ecological Application

CityNet produced realistic patterns of biotic and anthropogenic acoustic activity in the urban soundscape at two study sites of low and high urban intensity (Figure 2B and C). At both sites, biotic acoustic activity peaked just after sunrise and declined rapidly after sunset. A second peak of biotic acoustic activity was recorded at sunset at the low urban intensity site but not at the high urban intensity site. At both sites anthropogenic acoustic activity rose sharply after sunrise, remained constant throughout the day and declined after sunset.

## DISCUSSION

Both CityBioNet and CityAnthroNet outperformed the competing algorithms on the CitySound2017_*test*_ dataset. CityBioNet performed better than bulbul on noisy recordings from the urban environment; it was robust to more non-biotic sounds, including road traffic, air traffic and rain. Being robust to the sound of road traffic supports the suitability of CityBioNet for use in cities, as the urban soundscape is dominated by the sound of road traffic (Fairbrass *et al.* 2017) which has been shown to bias several of the AIs tested here (Fuller *et al.* 2015; Fairbrass *et al.* 2017). The sound of rain has also been shown to bias several AIs (Depraetere *et al.* 2012; Gasc *et al.* 2015b; Fairbrass *et al.* 2017) and the development of a method that is robust to this sound is a considerable contribution to the field of ecoacoustics. The urban biotic soundscape is dominated by the sounds emitted by birds (Fairbrass *et al.* 2017), and the good performance of bulbul, an algorithm for measuring exclusively bird sounds, on the CitySounds2017_*test*_ dataset, confirms this. Birds are used as indicator species in existing urban biodiversity monitoring schemes (Kohsaka *et al.* 2013) using data collected from traditional forms of biodiversity survey. The algorithms developed here could be used to support such existing schemes by making it easier to collect data on these indicator taxa.

CityNet is the only method currently available for measuring both biotic and anthropogenic acoustic activity using a single system in noisy audio data from urban environments. There is increasing evidence that anthropogenic noise affects wildlife in a variety of ways including altering communication behaviour (Gil & Brumm 2014) and habitat use (Deichmann *et al.* 2017). However, these investigations are limited in scale by the use of resource intensive methods of measuring biotic and anthropogenic sound in the environment or from audio data. Others rely on AIs (Pieretti & Farina 2013) which have been shown to be unreliable in acoustically disturbed environments (Fairbrass *et al.* 2017). CityNet could facilitate the investigation of the impacts of anthropogenic activities on wildlife populations at scales not currently possible with traditional acoustic analysis methods.

CityBioNet clearly outperformed all the AIs tested, but the difference in performance between CityAnthroNet and the competing algorithm for measuring anthropogenic acoustic activity (NDSI_*anthro*_) was much less marked. These results suggest that the measurement of biotic sound in noisy audio data from urban environments requires more sophisticated algorithms than the measurement of anthropogenic sound. Possibly anthropogenic sounds are more easily separable from other sounds in frequency space, a theory which is the basis of a number of AIs (Boelman *et al.* 2007; Kasten *et al.* 2012), facilitating the use of human defined algorithms such as NDSI_*anthro*_. Whereas, because biotic sounds occur in a frequency space shared with anthropogenic and geophonic sounds (Fairbrass *et al.* 2017), algorithms such as AIs which only use a small number of features to discriminate sounds are not sufficient for use in cities. Therefore, ML algorithms which are able to utilise larger numbers of features to discriminate sounds, such as the CNNs implemented in the CityNet system, are better able to detect biotic sounds in recordings that also contain non-biotic sounds. A recent unsupervised method developed by Lin, Fang and Tsao (2017) to separate biological sounds from long recordings could be used as a pre-processing step to further improve CityNet’s performance.

Low cost acoustic sensors and algorithms for the automatic measurement of biotic sound in audio data is facilitating the assessment and monitoring of biodiversity at large temporal and spatial scales (Sueur & Farina 2015), but to date this technology has only been deployed in non-urban environments (e.g. Aide *et al.* 2013). In cities, the availability of mains power and Wifi connections is supporting the development of the urban Internet of Things (IoT) using sensors integrated into existing infrastructure to monitor environmental factors including air pollution, noise levels, and energy use (Zanella *et al.* 2014). The CityNet system could be integrated into an IoT sensing network to facilitate large-scale urban environmental assessment. Large-scale deployment of algorithms such as CityNet requires low power usage and fast running times. One way to help to achieve this aim would be to combine the two networks (CityBioNet and CityAnthroNet) into one CNN which predicts both biotic and anthropogenic acoustic activity simultaneously.

An expansion of CityNet to ultrasonic frequencies would increase the generality of the tool as it could be used to monitor species in cities that emit sounds at frequencies higher than 12 kHz such as bats and some invertebrates. Bats are frequently used as ecological indicators because they are sensitive to environmental changes (Walters *et al.* 2013). Acoustic methods are commonly used to monitor bat populations using passive ultrasonic recorders meaning bat researchers and conservationists are faced with the challenge of extracting meaningful information from large volumes of audio data. The development of automated methods for measuring bat calls in ultrasonic data has focused to date on the identification of bat species calls and many algorithms are proprietary (e.g., Szewczak 2010; Wildlife Acoustics 2017). The development of an open-source algorithm that produces community-level measures of bats would be a valuable addition to the toolbox of bat researchers and conservationists.

Retraining CityNet with labelled audio data from other cities would make it possible to use the system to monitor urban biotic and anthropogenic acoustic activity more widely. However, as London is a large and heterogeneous city, CityNet has been trained using a dataset containing sounds that characterise a wide range of urban environments. Our data collection was restricted to a single week at each study site, which limits our ability to assess the ability of CityNet system to detect environmental changes. Future work should focus on the collection of longitudinal acoustic data to assess the sensitivity of the algorithms to detect environmental changes. Our use of human labellers would have introduced subjectivity and bias into our dataset. The task of annotating large audio datasets from acoustically complex urban environments is highly resource intensive, a problem which has been recently tackled with citizen scientists to create the UrbanSounds and UrbanSound8k datasets using audio data from New York city, USA (Salamon, Jacoby & Bello 2014). These comprise short snippets of 10 different urban sounds such as jackhammers, engines idling and gunshots. These datasets do not fully represent the characteristics of urban soundscapes for three reasons. Firstly, they assume only one class of sound is present at each time, while in fact multiple sound types can be present at one time (consider a bird singing while an aeroplane flies overhead). Secondly, they only include anthropogenic sounds, while CityNet measures both anthropogenic and biotic sounds. Finally, each file in these datasets has a sound present, while urban soundscapes contain many periods of silence or geophonic sounds, two important states which are not present in UrbanSounds and UrbanSounds8k. Due to these factors, these datasets are unsuitable for the purpose of this research project, although recent work has overcome a few of these shortcoming using synthesised soundscape data (Salamon *et al.* 2017). This highlights the need for an internationally coordinated effort to create a consistently labelled audio dataset from cities to support the development of automated urban environmental assessment systems with international application.

## Conclusions

The CityNet system for measuring biotic and anthropogenic acoustic activity in noisy urban audio data outperformed the state-of-the-art algorithms for measuring biotic and anthropogenic sound in entire audio recordings. Integrated into an IoT network for recording and analysing audio data in cities it could facilitate urban environmental assessment at greater scales than has been possible to date using traditional methods of biodiversity assessment.

We make our system available open source in combination with two expertly annotated urban soundscape datasets to facilitate future research development in this field.

## AUTHOR CONTRIBUTION STATEMENT

AF, MF, HT and KJ conceived ideas and designed methodology; AF collected the data; AF and MF analysed the data and led the writing of the manuscript. All authors contributed critically to the drafts and gave final approval for publication.

## ACKNOWLEDGMENTS

We thank multiple site owners and managers for supporting the study by providing access to recording sites, and multiple acoustic annotators and a transport expert for help creating the CitySounds2017 dataset. We were financially supported by a BHP Billiton Sustainable Resources for Sustainable Cities Catalyst Grant and by the Engineering and Physical Sciences Research Council (EPSRC) through a doctoral training grant (EP/G037698/1) to H.T., and EPSRC grant (EP/K015664/1) to K.E.J, G.B. and M.F.

## DATA ACCESSIBILITY

All recordings and annotations in the CitySounds2017 dataset and all Python code underlying the CityNet algorithms are available on Figshare (https://figshare.com/s/adab62c0591afaeafedd).

## SUPPORTING INFORMATION

### Section S1: Supplementary Methods

#### Normalisation Methods

The four normalisation methods used are as follows:

1. The entire spectrogram *S* was subtracted from each row in *W*_*s*_. This helped to act as a noise-reducing normalisation strategy
2. Each row of *W*_*s*_ was whitened to have zero mean and unit variance.
3. Each value in *W*_*s*_ was whitened to have zero mean and unit variance.
4. Each value in *W*_*s*_ was divided by the maximum value in *W*_*s*_.

#### Prediction Process

Both CityBioNet and CityAnthroNet have a convolutional layer with 32 filters, followed by a max pooling layer, then another 32-filter convolutional layer and finally two dense layers (with 128 units) before a binary class output - see Figure 1 for an overview of the network architecture. For nonlinearities very leaky rectifiers were used (Maas, Hannun & Ng 2013), and Dropout (Srivastava *et al.* 2014) was used to help to regularise the network and batch normalisation (Ioffe & Szegedy 2015) to increase the speed of convergence during training. The network was trained for 30 epochs using the Adam (Kingma & Ba 2015) update scheme with a learning rate of 0.0005. An ensemble of five such networks was trained using the same architecture and training data, but with different random initialisations. The final predictions are made by averaging together the predictions of each member in the ensemble.

**Table S1.** Details of acoustic recording sites across Greater London, UK. Sites separated into two groups illustrating whether recordings from sites were included in the CitySounds2017_*train*_ or CitySounds2017_*test*_ datasets. Urban intensity categories defined based on the predominant land cover surrounding sites within a 500m radius: (i) high (contiguous multi-storey buildings); (ii) medium (detached and semi-detached housing); and (iii) low (fields and/or woodland). DD denotes decimal degrees. In terms of site type, C denotes church or churchyard, CG denoted community garden, GR denotes green roof, GW denotes green wall, and NR denotes nature reserve.

| Site Code | Site Type | Survey<br>Start Date | Survey<br>End Date | Latitude<br>(DD) | Longitude<br>(DD) | Urban<br>Intensity |
| --- | --- | --- | --- | --- | --- | --- |
| **CitySounds2017<sub>train</sub>** |  |  |  |  |  |  |
| RM14 3YB | C | 11/06/2013 | 19/06/2013 | 51.55121 | 0.266853 | Low |
| W8 4LA | C | 21/06/2013 | 28/06/2013 | 51.50223 | -0.19147 | High |
| SW15 4LA | C | 02/07/2013 | 07/07/2013 | 51.44914 | -0.23697 | Medium |
| NW1 | C | 24/06/2013 | 01/07/2013 | 51.5105 | -0.20574 | High |
| SW11 2PN | C | 16/08/2013 | 23/08/2013 | 51.47057 | -0.16973 | High |
| E4 7EN | C | 06/10/2013 | 13/10/2013 | 51.63101 | 0.001266 | High |
| SE1 2RT 7 | GR | 19/05/2014 | 27/05/2014 | 51.30.16N | 0.4.53W | High |
| SE1 2RT 10 | GR | 19/05/2014 | 27/05/2014 | 51.30.16N | 0.4.50W | High |
| SW1W 0QP | GW | 30/05/2014 | 06/06/2014 | 51.49627 | -0.14489 | High |
| SW1E 6BN | GR | 30/05/2014 | 06/06/2014 | 51.4981 | -0.14138 | High |
| SE11 6DN | GR | 11/06/2014 | 20/06/2014 | 51.49313 | -0.11199 | High |
| SE4 1SA | GR | 20/06/2014 | 30/06/2014 | 51.45817 | -0.02751 | Medium |
| WC2N 6RH | GR | 01/07/2014 | 10/07/2014 | 51.50706 | -0.12388 | High |
| CR0 1SG | C | 02/07/2014 | 09/07/2014 | 51.3722 | -0.10604 | High |
| CR0 | C | 02/07/2014 | 09/07/2014 | 51.33934 | -0.01266 | Medium |
| RM2 5EL | C | 10/07/2014 | 17/07/2014 | 51.58773 | 0.201817 | Medium |
| RM4 1LD | C | 10/07/2014 | 17/07/2014 | 51.62349 | 0.223904 | Low |
| SE22 0SD | GR | 28/07/2014 | 04/08/2014 | 51.45332 | -0.05583 | Medium |
| TW7 6BE | C | 30/07/2014 | 06/08/2014 | 51.4719 | -0.31981 | Medium |
| W4 2PH | C | 30/07/2014 | 06/08/2014 | 51.48308 | -0.25326 | Medium |
| SE6 | C | 19/08/2014 | 26/08/2014 | 51.42804 | -0.01095 | Medium |
| SE8 4EA | C | 19/08/2014 | 27/08/2014 | 51.46841 | -0.02344 | Medium |
| IG11 0FJ | GR | 21/08/2014 | 01/09/2014 | 51.52069 | 0.109187 | Medium |
| W5 5EQ | GR | 28/08/2014 | 05/09/2014 | 51.50975 | -0.30812 | Medium |
| E14 0EY | C | 02/09/2014 | 10/09/2014 | 51.51072 | -0.01192 | High |
| E1 0NR | C | 03/09/2014 | 11/09/2014 | 51.51676 | -0.04122 | Medium |
| SE10 9EY | GR | 05/09/2014 | 12/09/2014 | 51.4849 | 0.006003 | Medium |
| N2 9BX | GR | 15/09/2014 | 22/09/2014 | 51.59274 | -0.16569 | Medium |
| SW6 6DU | GR | 16/09/2014 | 23/09/2014 | 51.47369 | -0.21695 | Medium |
| SE6 4PL | CG | 24/05/2015 | 01/06/2015 | 51.43821 | -0.02711 | Medium |
| W1T 4BQ | GR | 22/06/2015 | 30/06/2015 | 51.52143 | -0.13836 | High |
| N4 1ES | NR | 23/06/2015 | 02/07/2015 | 51.57656 | -0.1017 | Medium |
| TN14 7QB | NR | 25/06/2015 | 03/07/2015 | 51.31364 | 0.067323 | Low |
| NW3 3RY | NR | 14/07/2015 | 22/07/2015 | 51.54357 | -0.16054 | High |
| N8 8JD | CG | 11/07/2015 | 19/07/2015 | 51.58333 | -0.13292 | Medium |
| KT18 6AP | NR | 27/07/2015 | 05/08/2015 | 51.29036 | -0.26158 | Low |
| NW2 3SH | NR | 11/08/2015 | 18/08/2015 | 51.55287 | -0.20628 | Medium |
| N17 | CG | 17/08/2015 | 27/08/2015 | 51.59105 | -0.0549 | High |
| RM4 1PL | C | 27/08/2015 | 04/09/2015 | 51.61588 | 0.18189 | Medium |
| SE23 2NZ | NR | 16/09/2015 | 23/09/2015 | 51.43224 | -0.05197 | Medium |
| NW3 2BZ | NR | 17/09/2015 | 25/09/2015 | 51.55181 | -0.16259 | Medium |
| NW1 0TA | NR | 15/10/2015 | 22/10/2015 | 51.54073 | -0.13613 | High |
| SE15 4EE | CG | 13/10/2015 | 20/10/2015 | 51.46301 | -0.07519 | Medium |
| RM15 4HX | NR | 20/10/2015 | 28/10/2015 | 51.51749 | 0.261494 | Low |
| **CitySounds2017<sub>test</sub>** |  |  |  |  |  |  |
| W11 2NN | C | 08/07/2013 | 16/07/2013 | 51.53452 | -0.12957 | High |
| WC2H 8LG | C | 08/07/2013 | 14/07/2013 | 51.51521 | -0.12823 | High |
| HA8 6RB | C | 23/07/2013 | 30/07/2013 | 51.60862 | -0.2899 | Medium |
| HA5 3AA | C | 23/07/2013 | 30/07/2013 | 51.59478 | -0.37885 | Medium |
| SE23 | C | 06/09/2013 | 13/09/2013 | 51.45047 | -0.05146 | Medium |
| SE3 | C | 06/09/2013 | 13/09/2013 | 51.46261 | 0.001164 | Medium |
| CR8 | C | 15/09/2013 | 22/09/2013 | 51.3305 | -0.09394 | Medium |
| CR0 5EF | C | 15/09/2013 | 22/09/2013 | 51.37199 | -0.05031 | Medium |
| E10 5JP | C | 06/10/2013 | 13/10/2013 | 51.56386 | -0.01604 | Medium |
| SW15 4JY | GR | 27/08/2014 | 03/09/2014 | 51.45012 | -0.23859 | Medium |
| IG6 2XL | CG | 08/05/2015 | 15/05/2015 | 51.60046 | 0.095681 | Low |
| E2 9RR | NR | 25/05/2015 | 02/06/2015 | 51.5295 | -0.05875 | High |
| TW7 6ER | C | 23/06/2015 | 30/06/2015 | 51.46711 | -0.3454 | Medium |
| BR2 0EG | C | 17/07/2015 | 26/07/2015 | 51.4047 | 0.012974 | Medium |
| BR2 8LB | C | 31/07/2015 | 07/08/2015 | 51.38029 | 0.042746 | Medium |
| BR6 7US | C | 31/07/2015 | 07/08/2015 | 51.33605 | 0.054201 | Low |
| BR4 | C | 18/08/2015 | 25/08/2015 | 51.38261 | -0.00868 | Medium |
| DA5 | NR | 24/08/2015 | 01/09/2015 | 51.42268 | 0.156502 | Medium |
| CM16 7NP | NR | 08/09/2015 | 15/09/2015 | 51.65396 | 0.101227 | Low |

**Figure S1.**
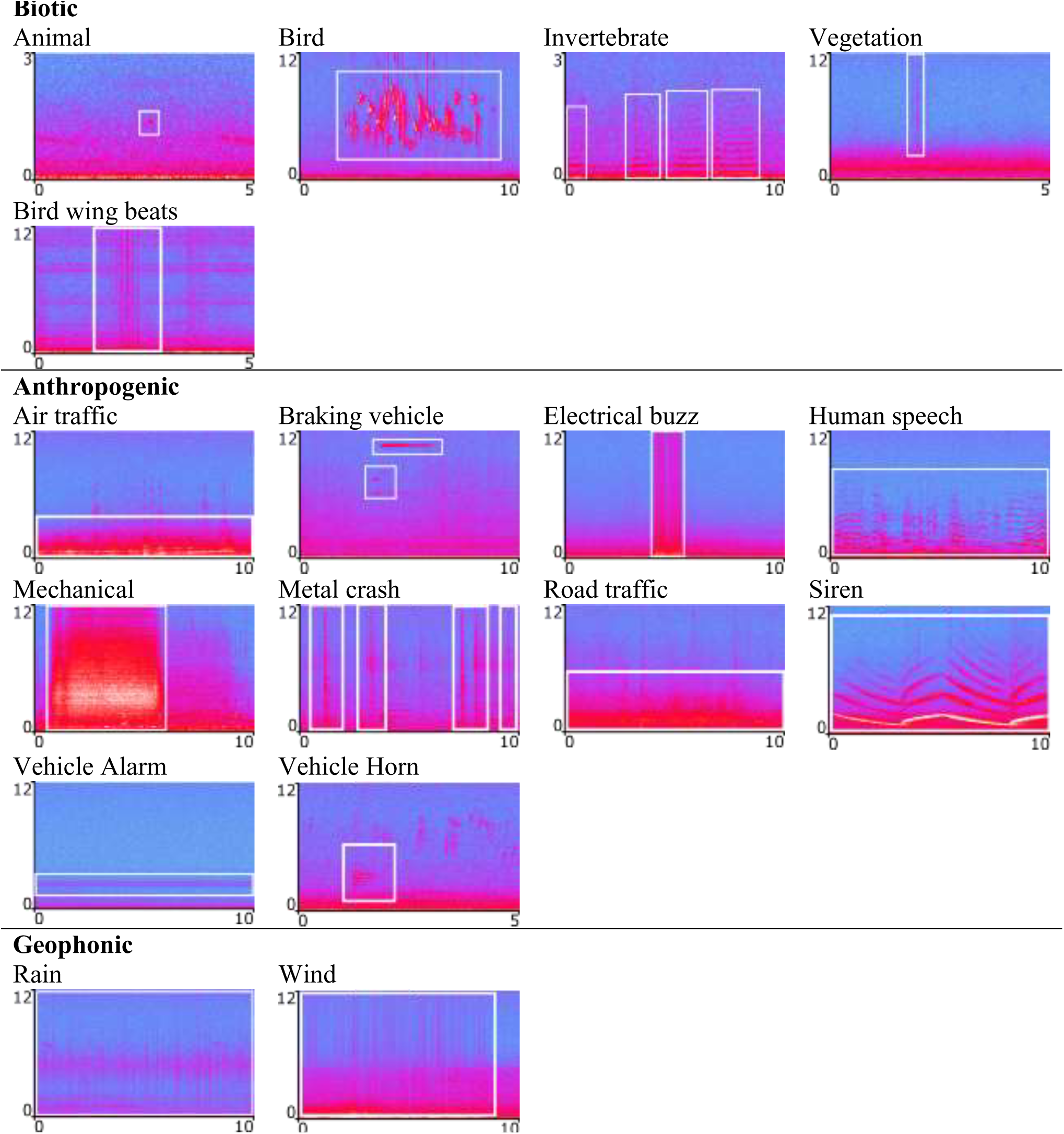
Examples of all sound types present in CitySounds2017. ‘Animal’ denotes biotic sounds that could not be taxonomically identified. Unidentified sounds not shown due to wide range of sound types within this group. Data is represented in spectrograms (FFT nonoverlapping Hamming window size=1024) where blue to yellow corresponds to sound amplitude (dB). Frequency (kHz) and time (s) are represented on the y‐ and x-axes, respectively. Spectrograms represent biotic (sounds generated by non-human biotic organisms), anthropogenic (sounds associated with human activities including human speech) and geophonic sounds.

## REFERENCES

Acevedo, M.A., Corrada-Bravo, C.J., Corrada-Bravo, H., Villanueva-Rivera, L.J. & Aide, T.M. (2009) Automated classification of bird and amphibian calls using machine learning: A comparison of methods. Ecological Informatics, 4, 206–214.

Aide, T.M., Corrada-Bravo, C., Campos-Cerqueira, M., Milan, C., Vega, G. & Alvarez, R. (2013) Real-time bioacoustics monitoring and automated species identification. 1, e103. Available: http://www.ncbi.nlm.nih.gov/pmc/articles/PMC3719130/pdf/peerj-01-103.pdf Accessed: 19/12/2016

Aronson, M.F.J., La Sorte, F.A., Nilon, C.H., Katti, M., Goddard, M.A., Lepczyk, C.A., Warren, P.S., Williams, N.S.G., Cilliers, S., Clarkson, B., Dobbs, C., Dolan, R., Hedblom, M., Klotz, S., Kooijmans, J.L., Kühn, I., MacGregor-Fors, I., McDonnell, M., Mörtberg, U., Pyšek, P., Siebert, S., Sushinsky, J., Werner, P. & Winter, M. (2014) A global analysis of the impacts of urbanization on bird and plant diversity reveals key anthropogenic drivers. 281, 20133330. Available: http://rspb.royalsocietypublishing.org/content/281/1780/20133330.abstract Accessed: 02/12/2016

Beninde, J., Veith, M. & Hochkirch, A. (2015) Biodiversity in cities needs space: a metaanalysis of factors determining intra-urban biodiversity variation. Ecology Letters, 18, 581–592.

Boelman, N.T., Asner, G.P., Hart, P.J. & Martin, R.E. (2007) Multi-trophic invasion resistance in Hawaii: Bioacoustics, field surveys, and airborne remote sensing. Ecological Applications, 17, 2137–2144.

Chesmore, E. & Ohya, E. (2004) Automated identification of field-recorded songs of four British grasshoppers using bioacoustic signal recognition. Bulletin of Entomological Research, 94, 319–330.

Cohen, J. (1992) Statistical power analysis. Current directions in psychological science, 1, 98–101.

Crouse, D.L., Pinault, L., Balram, A., Hystad, P., Peters, P.A., Chen, H., van Donkelaar, A., Martin, R.V., Ménard, R., Robichaud, A. & Villeneuve, P.J. (2017) Urban greenness and mortality in Canada's largest cities: a national cohort study. The Lancet Planetary Health, 1, e289–e297.

Deichmann, J.L., Hernández-Serna, A., Delgado C, J.A., Campos-Cerqueira, M. & Aide, T.M. (2017) Soundscape analysis and acoustic monitoring document impacts of natural gas exploration on biodiversity in a tropical forest. Ecological Indicators, 74, 39–48.

Depraetere, M., Pavoine, S., Jiguet, F., Gasc, A., Duvail, S. & Sueur, J. (2012) Monitoring animal diversity using acoustic indices: implementation in a temperate woodland. Ecological Indicators, 13, 46–54.

Dieleman, S., Schlüter, J., Raffel, C., Olson, E., Sønderby, S.K., Nouri, D., Maturana, D., Thoma, M., Battenberg, E., Kelly, J., De Fauw, J., Heilman, M., de Almeida, D.M., McFee, B., Weideman, H., Takács, G., de Rivaz, P., Crall, J., Sanders, G., Rasul, K., Liu, C., French, G. & Degrave, J. (2015) Lasagne. Available: http://dx.doi.org/10.5281/zenodo.27878 Accessed: 19/09/2017

Digby, A., Towsey, M., Bell, B.D. & Teal, P.D. (2013) A practical comparison of manual and autonomous methods for acoustic monitoring. Methods in Ecology and Evolution, 4, 675–683.

Faeth, S.H., Bang, C. & Saari, S. (2011) Urban biodiversity: patterns and mechanisms. Year in Ecology and Conservation Biology, 1223, 69–81.

Fairbrass, A.J., Rennett, P., Williams, C., Titheridge, H. & Jones, K.E. (2017) Biases of acoustic indices measuring biodiversity in urban areas. Ecological Indicators, 83, 169–177.

Farinha-Marques, P., Lameiras, J., Fernandes, C., Silva, S. & Guilherme, F. (2011) Urban biodiversity: a review of current concepts and contributions to multidisciplinary approaches. Innovation: The European Journal of Social Science Research, 24, 247–271.

Fuller, S., Axel, A.C., Tucker, D. & Gage, S.H. (2015) Connecting soundscape to landscape: Which acoustic index best describes landscape configuration? Ecological Indicators, 58, 7.07–7.15

Gasc, A., Pavoine, S., Lellouch, L., Grandcolas, P. & Sueur, J. (2015a) Acoustic indices for biodiversity assessments: Analyses of bias based on simulated bird assemblages and recommendations for field surveys. Biological Conservation, 191, 306–312.

Gasc, A., Pavoine, S., Lellouch, L., Grandcolas, P. & Sueur, J. (2015b) Acoustic indices for biodiversity assessments: Analyses of bias based on simulated bird assemblages and recommendations for field surveys. Biological Conservation, 191, 306–312.

Gil, D. & Brumm, H. (2014) Acoustic communication in the urban environment: patterns, mechanisms, and potential consequences of avian song adjustments. Avian urban ecology (eds D. Gil & H. Brumm), pp. 69–83. Oxford University Press, Oxford, UK.

Grill, T. & Schlüter, J. (2017) Two Convolutional Neural Networks for Bird Detection in Audio Signals. 25th European Signal Processing Conference (EUSIPCO2017). Kos, Greece.

Hall, D.M., Camilo, G.R., Tonietto, R.K., Smith, D.H., Ollerton, J., Ahrné, K., Arduser, M., Ascher, J.S., Baldock, K.C. & Fowler, R. (2016) The city as a refuge for insect pollinators. Conservation Biology, 31, 24–29.

Hunter, J.D. (2007) Matplotlib: A 2D graphics environment. Computing In Science & Engineering, 9, 90–95.

Ioffe, S. & Szegedy, C. (2015) Batch normalization: Accelerating deep network training by reducing internal covariate shift. Proceedings of the 32nd International Conference on Machine Learning, pp. 448–456. Lille, France.

Kasten, E.P., Gage, S.H., Fox, J. & Joo, W. (2012) The remote environmental assessment laboratory's acoustic library: An archive for studying soundscape ecology. Ecological Informatics, 12, 50–67.

Kingma, D. & Ba, J. (2015) Adam: A Method for Stochastic Optimization. Proceedings of the International Conference on Learning Representations 2015. San Deigo, USA.

Kohsaka, R., Pereira, H.M., Elmqvist, T., Chan, L., Moreno-Peñaranda, R., Morimoto, Y., Inoue, T., Iwata, M., Nishi, M. & da Luz Mathias, M. (2013) Indicators for management of urban biodiversity and ecosystem services: city biodiversity index. Urbanization, biodiversity and ecosystem services: challenges and opportunities (eds T. Elmqvist, M. Fragkias, J. Goodness, B. Güneralp, P.J. Marcotullio, R.I. McDonald, S. Parnell, M. Schewenius, M. Sendstad, K.C. Seto & C. Wilkinson), pp. 699–718. Springer, Netherlands.

LeCun, Y., Bengio, Y. & Hinton, G. (2015) Deep learning. Nature, 521, 436–444.

Lee, H., Pham, P., Largman, Y. & Ng, A.Y. (2009) Unsupervised feature learning for audio classification using convolutional deep belief networks. Proceedings of the 22nd International Conference on Neural Information Processing Systems, pp. 1096–1104. Istanbul, Turkey.

Lin, T.-H., Fang, S.-H. & Tsao, Y. (2017) Improving biodiversity assessment via unsupervised separation of biological sounds from long-duration recordings. 7. Available: https://www.nature.com/articles/s41598-017-04790-7 Accessed: 19/09/2017

Maas, A.L., Hannun, A.Y. & Ng, A.Y. (2013) Rectifier nonlinearities improve neural network acoustic models. Proceedings of the 30th International Conference on Machine Learning. Atlanta, USA.

McFee, B., Raffel, C., Liang, D., Ellis, D.P., McVicar, M., Battenberg, E. & Nieto, O. (2015) librosa: Audio and music signal analysis in python. Proceedings of the 14th python in science conference, pp. 18–25. Austin, Texas.

Natural England (2016) Links between natural environments and mental health: evidence briefing. Available: http://publications.naturalengland.org.uk Accessed: 24/11/2017

Pedregosa, F., Varoquaux, G., Gramfort, A., Michel, V., Thirion, B., Grisel, O., Blondel, M., Prettenhofer, P., Weiss, R., Dubourg, V., Vanderplas, J., Passos, A., Cournapeau, D., Brucher, M. & Perrot, M.D., E. (2011) Scikit-learn: Machine Learning in Python. Journal of machine learning research, 12, 2825–2830.

Pieretti, N. & Farina, A. (2013) Application of a recently introduced index for acoustic complexity to an avian soundscape with traffic noise. The Journal of the Acoustical Society of America, 134, 891–900.

Pieretti, N., Farina, A. & Morri, D. (2011) A new methodology to infer the singing activity of an avian community: the Acoustic Complexity Index (ACI). Ecological Indicators, 11, 868–873.

Pijanowski, B.C., Villanueva-Rivera, L.J., Dumyahn, S.L., Farina, A., Krause, B.L., Napoletano, B.M., Gage, S.H. & Pieretti, N. (2011) Soundscape ecology: the science of sound in the landscape. Bioscience, 61, 203–216.

Python Software Foundation (2016) Python Language Reference. Available: http://www.python.org Accessed: 19/09/2017

R Core Team (2017) R: A language and environment for statistical computing. Available: http://www.R-project.org Accessed: 31/10/2014

Salamon, J., Jacoby, C. & Bello, J.P. (2014) A dataset and taxonomy for urban sound research. ACMMM’14, pp. 1041–1044. Association for Computing Machinery, Orlando, USA.

Salamon, J., MacConnell, D., Cartwright, M., Li, P. & Bello, J.P. (2017) Scaper: A library for soundscape synthesis and augmentation. 2017 IEEE Workshop on Applications of Signal Processing to Audio and Acoustics. New Paltz, NY.

Srivastava, N., Hinton, G.E., Krizhevsky, A., Sutskever, I. & Salakhutdinov, R. (2014) Dropout: a simple way to prevent neural networks from overfitting. Journal of machine learning research, 15, 1929–1958.

Stowell, D. & Plumbley, M.D. (2014) Automatic large-scale classification of bird sounds is strongly improved by unsupervised feature learning. 2, e488. Available: http://dx.doi.org/10.7717/peerj.488 Accessed: 09/12/2016

Stowell, D., Wood, M., Stylianou, Y. & Glotin, H. (2016) Bird detection in audio: a survey and a challenge. 2016 IEEE 26th International Workshop on Machine Learning for Signal Processing, pp. 1–6. IEEE, Vietri sul Mare, Italy.

Sueur, J., Aubin, T. & Simonis, C. (2008) Equipment review: seewave, a free modular tool for sound analysis and synthesis. Bioacoustics, 18, 213–226.

Sueur, J. & Farina, A. (2015) Ecoacoustics: the Ecological Investigation and Interpretation of Environmental Sound. Biosemiotics, 8, 493–502.

Sueur, J., Farina, A., Gasc, A., Pieretti, N. & Pavoine, S. (2014) Acoustic Indices for Biodiversity Assessment and Landscape Investigation. Acta Acustica united with Acustica, 100, 772–781.

Szewczak, J.M. (2010) SonoBat. Available: www.sonobat.com Accessed: 29/05/2014

The Theano Development Team, Al-Rfou, R., Alain, G., Almahairi, A., Angermueller, C., Bahdanau, D., Ballas, N., Bastien, F., Bayer, J. & Belikov, A. (2016) Theano: A Python framework for fast computation of mathematical expressions. Available: https://arxiv.org/abs/1605.02688 Accessed: 19/09/2017

Towsey, M., Wimmer, J., Williamson, I. & Roe, P. (2014) The Use of Acoustic Indices to Determine Avian Species Richness in Audio-recordings of the Environment. Ecological Informatics, 21, 110–119.

UN-DESA (2016) The World's Cities in 2016. Data Booklet. Available: http://www.un.org/en/development/desa/population/ Accessed: 10/02/2017

Villanueva-Rivera, L.J. & Pijanowski, B.C. (2014) Package ‘soundecology’. Soundscape ecology. Available: http://cran.r-project.org/web/packages/soundecology/index.html Accessed: 15/04/2015

Villanueva-Rivera, L.J., Pijanowski, B.C., Doucette, J. & Pekin, B. (2011) A primer of acoustic analysis for landscape ecologists. Landscape Ecology, 26, 1233–1246.

Walters, C.L., Collen, A., Lucas, T., Mroz, K., Sayer, C.A. & Jones, K.E. (2013) Challenges of Using Bioacoustics to Globally Monitor Bats. Bat Evolution, Ecology, and Conservation, pp. 479–499. Springer.

Walters, C.L., Freeman, R., Collen, A., Dietz, C., Brock Fenton, M., Jones, G., Obrist, M.K., Puechmaille, S.J., Sattler, T., Siemers, B.M., Parsons, S. & Jones, K.E. (2012) A continental-scale tool for acoustic identification of European bats. Journal of Applied Ecology, 49, 1064–1074.

Wildlife Acoustics, I. (2017) Kaleidoscope Analysis Software. Available: https://www.wildlifeacoustics.com/products/kaleidoscope-software-ultrasonic Accessed: 24/08/2017

Zamora-Gutierrez, V., Lopez-Gonzalez, C., MacSwiney Gonzalez, M.C., Fenton, B., Jones, G., Kalko, E.K., Puechmaille, S.J., Stathopoulos, V. & Jones, K.E. (2016) Acoustic identification of Mexican bats based on taxonomic and ecological constraints on call design. Methods in Ecology and Evolution, 7, 1082–1091.

Zanella, A., Bui, N., Castellani, A., Vangelista, L. & Zorzi, M. (2014) Internet of things for smart cities. IEEE Internet of Things journal, 1, 22–32.

